# Circulating MicroRNAs indicative of sex and stress in the European seabass (*Dicentrarchus labrax*): toward the identification of new biomarkers

**DOI:** 10.1101/2023.07.03.547501

**Authors:** Camille Houdelet, Eva Blondeau-Bidet, Mathilde Estevez-Villar, Xavier Mialhe, Sophie Hermet, François Ruelle, Gilbert Dutto, Aline Bajek, Julien Bobe, Benjamin Geffroy

## Abstract

MicroRNAs (miRNAs) constitute a new category of biomarkers. Studies on miRNAs in non-mammalian species have drastically increased in the last few years. Here, we explored the use of miRNAs as potential, poorly-invasive markers, to identify sex and characterize acute stress in fish. The European seabass (*Dicentrarchus labrax*) was chosen as model because of its rapid response to stress and its specific sex determination system, devoid of sexual chromosomes. We performed a small RNA-sequencing analysis in the blood plasma of males and females’ European seabass (mature and immature) as well as in the blood plasma of juveniles submitted to an acute stress and sampled throughout the recovery period (at 0h, 0.5h, 1.5h and 6h). In immature individuals, both miR-1388-3p and miR-7132a-5p were up-regulated in females, while miR-499a-5p was more abundant in males. However, no miRNAs were found to be differentially expressed between sexes in the blood plasma of mature individuals. For the acute stress analysis, five miRNAs (miR-155-5p, miR-200a-3p, miR-205-1-5p, miR-143-3p and miR-223-3p) followed cortisol production over time. All miRNAs identified were tested and validated by RT-qPCR on sequenced samples. A complementary analysis on the 3’UTR sequences of the European seabass allowed to predict potential mRNA targets, some of them being particularly relevant regarding stress regulation, e.g. the glucocorticoid and the mineralocorticoid receptor. The present study provides new avenues and recommendations on the use of miRNAs as biomarkers of sex or stress of the European seabass, with potential application on other fish species.

## 1. Introduction

Identifying new, poorly-invasive, technics to depict clear phenotypes is of major interest in fishery and aquaculture contexts (Brosset et al., 2021; Raposo de Magalhães et al., 2020). For instance, consumers are now highly concerned by the welfare of harvested fish, and accurately monitoring stress is thus becoming crucial for both industries. Managing sex-ratio of farmed population is also primordial for producers, since in many fish species i) there is a strong sexual size dimorphism, and a clear benefit in producing the fastest-growing sex and ii) having a sufficient number of males and females is essential to guarantee successful genetic programs. Sexing wild fish with poorly-invasive tools can also be valuable to fully understand population dynamics in marine animals (e.g., Tunas) in which it is impossible to distinguish males from females based on external phenotypic characteristics.

The European seabass (*Dicentrarchus labrax*) is a convenient model to scrutinize all the above-mentioned issues, as it brings together all questioning. First, it is a recognized model of interest in aquaculture and fisheries (Vandeputte et al., 2019). Second, reliable basal levels of cortisol (major indicator of stress) are highly difficult to obtain in this species (Sadoul et al., 2021; Vandeputte et al., 2016) and is considered very high in relation to other species (Samaras et al., 2016). Third, there is currently no practical tools to distinguish males from females in this species that possess a polygenic sex determination system, with effects of the environment (Geffroy et al., 2021; Vandeputte et al., 2007). Indeed, it is only possible to visually and accurately discriminate males from females based on their size when they reach a certain age (i.e., after 3 years; Chatain & Chavanne, 2009). Hence, having other, poorly-invasive tools, to monitor sex and stress would be a real asset for this species, but also for many other exhibiting similar characteristics. Circulating microRNAs (miRNAs) could well fulfill these roles.

MiRNAs are short and conserved non-coding sequences of nucleotides (20-22 nt) involved in the regulation of multiple biological processes by post-transcriptional repression of target mRNA (Bartel, 2018). MicroRNAs act as gene regulators through a base-pairing interaction with mRNAs, and a sequence similarity of 6 to 8 nucleotides (seed sequence) is sufficient to deregulate a mRNA. Thus, a given miRNA could, in theory, interact with multiple mRNAs (up to several hundreds), and one mRNA could be the target of multiple miRNA (Fabian et al., 2010). MiRNAs present very advantageous characteristics to track physiological states, so that they are considered as efficient biomarkers. First, miRNAs are highly conserved in evolution, which support the prominent hypothesis of a fundamental role in the biology of metazoan (Bartel, 2009; Desvignes et al., 2019; Wheeler et al., 2009). Second, they can be quantified in different tissues, but also in the extracellular part of various body fluids (e.g., serum, blood plasma, urine, saliva; Mohr and Mott 2015). Since their discovery, they have been widely used for health-associated diagnostics in humans, mainly for tumor detection (Duttagupta et al., 2011; Lu et al., 2005), but there is an increasing interest in using them as biomarkers in other species. MiRNAs were shown to display distinct levels of expression between the gonads of males and females in various fish species (Bhat et al., 2020; Gu et al., 2014; Jing et al., 2014; Tao et al., 2016). However, to the best of our knowledge, sex-specific miRNAs have not yet been described in fluids (like the plasma). A recent study conducted in rainbow trout (*Onchorynchus mykiss*) identified a repertoire of circulating miRNAs that reflect the physiological and reproductive state of the species (Cardona et al., 2021). Regarding stress, miRNAs were also shown to transduce different stressful environments (Raza et al., 2022), where the impact of xenobiotics (Burgos-Aceves et al., 2018), handling (Cadonic et al., 2020; Ikert et al., 2021) or thermal stress (Raza et al., 2022) have been pinpointed. Still, we do not yet know how circulating miRNAs correlate to stress-induced cortisol levels.

The main objectives of this study were to characterize the circulating miRNAs associated with sex and stress in the European seabass and to evaluate to what extent they can be easily quantified in the plasma by qPCR, with the ultimate goal of using them as biomarkers.

## 2. Material and methods

Experiments were performed in accordance with relevant guidelines and regulations provided by the ethic committee (no 36) of the French Ministry of Higher Education, Research and Innovation and the experiment received the following agreement number: APAFIS #30612-2021031812193539.

### 2.1. Sample collection

#### 2.1.1. Experiment 1: Sex identification

A total of forty immature (mean weight: 44 ± 11 g and length: 15 ± 1 cm) and twenty mature (1387 ± 303 g and 46 ± 4 cm) fish were randomly collected from the experimental aquaculture station of Ifremer (Palavas-les-Flots, France) and Aquanord aquaculture site (Gravelines, France) respectively. A blood sample (1 ml, see details below) was collected from each individual, as well as a piece of gonad for immature fish to histologically sex them (Fig. 1A). For qPCR analyses, supplementary individuals were collected and sexed (validation non-sequenced 1, Table 1) as well as fish from the stress experiment (validation non-sequenced 2, Table 1).

**Fig. 1.**
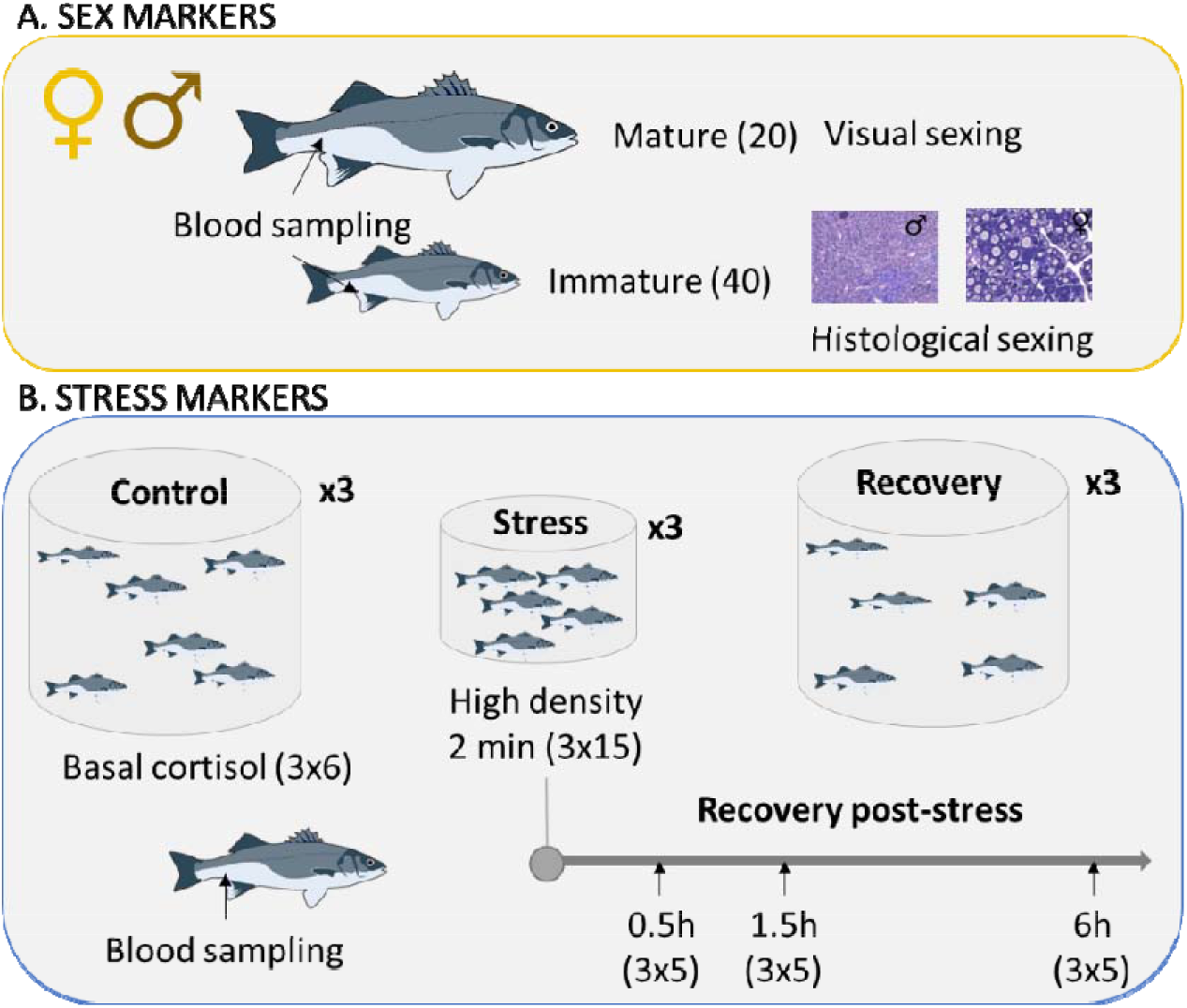
Summary of the two experiments conducted to identify (A) sex and (B) stress markers. The number in brackets represent the number of fish used

**Table 1.** Summary of the number of samples used for small RNA-seq and qPCR analyses

|  |  | Sex |  |  |  |
| --- | --- | --- | --- | --- | --- |
|  |  | small RNA-seq | qPCR | weight (g) | length (cm) |
| **Experiment 1: sex** | Male (Matures) | 3 |  | 1333 | 47 |
|  | Female (Matures) | 3 |  | 1281 | 45 |
|  | Male (Immatures) | 5 | 5 | 44 | 15 |
|  | Female (Immatures) | 5 | 5 | 41 | 15 |
| **Validation non-sequenced 1** | Male |  | 5 | 289 | 28 |
|  | Female |  | 5 | 398 | 30 |
| **Validation non-sequenced 2** | Male |  | 5 | 279 | 27 |
|  | Female |  | 4 | 484 | 32 |
| **Experiment 2: stress** | Male | 9 | 15 | 39 | 14 |
|  | Female | 11 | 12 | 45 | 15 |

#### 2.1.2. Experiment 2: Stress experiment

Juvenile European Seabass (n = 63, mean weight: 42.1 ± 15.7 g and length: 14.5 ± 1.5 cm) were maintained in a recirculating aquaculture system, with three tanks of 1.5 m^3^ each (21.0 ± 0.1 °C and water renewing at 1.2 m^3^/h), connected to a common biofilter tank. The three tanks were covered with a black tarpaulin ensuring they were visually isolated from the experimenters. The photoperiod (12L/12D) inside the tarpaulin was maintained using a specific a light system: AquaRay miniLED 500, 10000K white (Tropical Marine Center). Fish were fasted 24 h prior the start of the experiment. Beforehand the stress procedure, six fish per tank (n=18 in total) were quickly sampled and euthanized using benzocaine (150 mg/L). The blood was immediately collected by 3 experimenters, ensuring that all fish were collected within 2 minutes to avoid cortisol rise commonly reported after death (Sadoul and Geffroy, 2019). Thenceforward, 15 fish per tank were quickly placed in a 10L bucket at a density of 300kg/m3 for 2 minutes. Following this confinement challenge, they were placed in 3 recovering tanks (100L tanks, supplied with renewed water from the same original tank) to recover from the stress for 0.5, 1.5 and 6 hours (Fig. 1B). Supplementary individuals were collected for qPCR validation (n = 30) see details in Table 2).

**Table 2.** Summary of the number of samples used for small RNA-seq and qPCR validation of stress biomarkers

|  |  | Stress |  |  |  |
| --- | --- | --- | --- | --- | --- |
|  |  | small RNA-seq | qPCR | weight (g) | length (cm) |
| **Experiment stress** | **T0** | **4** | **12** | **32** | **13** |
|  | **T0.5** | **5** | **4** | **40** | **14** |
|  | **T1.5** | **5** | **8** | **36** | **14** |
|  | **T6** | **7** | **6** | **38** | **14** |

#### Blood sampling

At 0.5, 1.5 and 6 hours following the confinement challenge, a subsample of five fish was collected from each tank (total n=15 per recovering tank), euthanized with a lethal dose benzocaine (150mg/L) and sexed (note that one individual was not sexed). The blood was sampled from the caudal vein thanks to EDTA-coated syringes. After centrifugation (3000 x g, 5 min, 10°C) the plasma was allocated in two new tubes: one part for cortisol quantification (20µL) and the other part for small-RNA sequencing, stocked at –20°C and –80°C, respectively.

### 2.2. Identification of markers

#### 2.2.1 Histology

Gonads of immature European seabass were fixed in Bouin’s fluid for 6 to 8 h and rinsed in clear water for one hour. Then, they were rinsed in EtOH 70% for several days and placed in a dehydration automate (STP 120, MM, France). Each gonad was embedded in paraffin and cut at 5µm sections. Slides were stained using the Masson’s trichrome methods (MYREVA SS30, MM, France).

#### 2.2.2 Measurement of stress

Plasmatic cortisol was measured using a Cortisol ELISA kit (Neogen Lexington, KY, USA). According to the supplier, the cross-reactivity of the antibody with other steroids is as follows: prednisolone 47.5%, cortisone 15.7%, 11-deoxycortisol 15.0%, prednisone 7.83%, corticosterone 4.81%, 6_β_-hydroxycortisol 1.37%, 17-hydroxyprogesterone 1.36%. Following manufacturer’s instructions, samples or standard (Cortisol standard solution) were added in each well in duplicate and supplemented with the conjugated cortisol enzyme. After one hour of incubation, each well was washed and filled with the substrate. Absorbance was read at 650 nm with a microplate reader (Synergy HT, BioTek Instrument, VT, USA) after 30 minutes of incubation in the dark. To confirm the repeatability of the experiment, one sample was placed on the three different plates. Parallel displacement curves were obtained for plasma by comparing serial dilutions of pooled plasma (1:1 – 1:250) and the cortisol standard preparation (0.04 – 10 ng/ml). All values are expressed with the standard error.

#### 2.2.3 RNA extractions

Thawed plasma of each fish from the two experiments was homogenized in QIAzol lysis reagent (Beverly, MA, USA) following manufacturer’s instructions. The total RNA was resuspended in 15 µL of RNAse free water. MiRNAs were quantified using the smallRNA Analysis kit (DNF-470-0275) on a Fragment Analyzer (Agilent).

#### 2.2.4 Small RNA Sequencing

Following these steps, six matures (over the 20 collected) and ten immatures (over the 40 collected) individuals from first experiment (sex identification) and 21 individuals from the second experiment (stress experiment) were of sufficient quality and quantity, to be further processed (Table 1 and Table 2). Libraries were constructed using the NEXTFLEX Small RNA-seq kit v3 (Perkin Elmer, #NOVA-5132-05). Briefly, a 3’ Adenylated adapter was ligated to the 3’ end of 0,5 ng of microRNA and purified to remove 3’ adapter excess. A 5’ adapter was ligated to the 5’ end of the 3’ ligated microRNA. The resulting construction was purified to remove 5’ adapter excess. 5’ and 3’ ligated microRNAs underwent a reverse transcription using a M-MuLV reverse Transcriptase and a RT primer complementary to the 3’ adapter. Resulting cDNAs were used as a matrix in a 25 cycles PCR using a pair of uniquely barcoded primers. The resulting barcoded library was size selected on a Pippin HT (SAGE Science) using 3 % agarose cassette (#HTG3004) and aiming for a size range between 147 bp and 193 bp. Once size selected, libraries were verified on a Fragment Analyzer using the High Sensitivity NGS kit (#DNF-474-0500) and quantified using the KAPA Library quantification kit (Roche, ref. KK4824).

#### 2.2.5 MiRNA alignment and quantification

Image analyses and base calling were performed using the NovaSeq Control Software and Real-Time Analysis component (Illumina). Demultiplexing was performed using Illumina’s conversion software (bcl2fastq 2.20). The quality of the raw data was assessed using FastQC from the Babraham Institute and the Illumina software SAV (Sequencing Analysis Viewer).

The raw reads were trimmed using Cutadapt (version 3.5) (Martin, 2011) to remove the sequencing adapter (TGGAATTCTCGGGTGCCAAGG) at the 3′-end. Additionally, 4 bases were also trimmed from the 5′-end and 3′-end of the reads as indicated in the manual of NEXTflex Small RNA-Seq Kit v3 from Bio Scientific. Before counting step, samples with a rRNA degradation profile were filtered out. This resulted in 13 and 16 immature and mature fish, respectively, for the experiment on sex and 21 fish for the experiment on stress. MiRNA analysis was performed with Prost ! v0.7.60 pipeline (Desvignes et al., 2019). In this pipeline, alignment was performed with BBMap (Bushnell, 2014) version 38.90 to genome of interest: *Dicentrarchus labrax* GCA_000689215.1_seabass_V1.0. The miRNA annotation and counting were retrieved by Prost! (parameters in Text S1) from a custom annotation provided in miRBase v21 (available in *Prost!* Github) using *Gasterosteus aculeatus* miRNA sequences (the phylogenetically closest species).

### 2.3 Validation of markers by qPCR

#### 2.3.1. cDNA synthesis and quantitative real-time PCR

For the validation of markers by qPCR, total RNA from each of the non-sequenced samples (n = 56 individuals for sex and n = 30 individuals for stress; Table 1) were normalized at 10 ng/µL and the reverse transcription of RNA was done following the manufacturer instructions (miRCURY LNA RT Kit, Qiagen). We added 0.5µl of controls UniSp6 and cel-mir-39-3p to the samples as internal reference to check the efficiency of the reverse transcription and PCR amplification, respectively. The reverse transcription reaction was conducted on a total volume of 10 µL containing 5 µL of 2x miRCURY SYBR Green Master Mix, 1 µL of the resuspended primer mix (miRCURY LNA PCR Assay), 3µL of diluted cDNA and 1µL of RNase-free water. Quantitative RT-PCR was performed on a Light Cycler 480 System (Roche Life Science) with the following conditions: 95°C for 2 min, and 45 cycles of 95°C for 10sec, 56°C for 60sec.

In addition to the exogenous control (cel-miR-39-3p), we selected one endogenous miRNA based on the small RNA sequencing and that was stable regarding the different conditions: miR-23b-2-3p. Sequences of miRNAs primer are provided in Table S1.

#### 2.3.2 Statistical analysis

Differentially expressed miRNA were identified using one Bioconductor (Gentleman et al., 2004) package: DESeq2 1.32.0 (Love et al., 2014). Data were normalized using the default method for DESeq2 package.

MiRNA with adjusted p-value below 5% (according to the FDR method from Benjamini-Hochberg) were considered significantly differentially expressed between conditions: Sex (Male *vs* Female) or Stress (0 *vs* 0.5 *vs* 1.5 *vs* 6 hours).

For qPCR experiments, the differences between sex or timing after stress was assessed using the non-parametric Kruskal-Wallis test (Kruskal & Wallis, 1952). For RT-qPCR on non-sequenced samples for sexing (n = 46; Table 1), a linear mixed-effects model was applied to consider the experiment effect. To identify the miRNAs that presented a linear increase (or decrease) in expression over time, following the confinement challenge, we ran a loop in R to automatically detect all miRNAs (normalized counts) that are significantly correlated to the time. For cortisol level analyses, the differences between timing after stress was assessed using the non-parametric Kruskal-Wallis test. A Non-metric Multi-dimensional Scaling (NMDS) approach was used to compare individuals from the different time-points and that presented various cortisol levels. The package vegan v2.6-2 (Oksanen et al., 2013) was used and the construction of the dissimilarity matrix was based on the Bray-Curtis methods. All statistical analyses presented in this section were conducted with the R v 4.1.0 (Core Team, 2020).

### 2.4. Prediction of target

We first retrieved the 3’UTR sequences from the European seabass genome browser (http://seabass.mpipz.mpg.de/) that is based on the published genome of the European seabass (Tine et al., 2014). To identify miRNA targets, we used the freely available Perl script of TargetScanHuman 8.0. (https://www.targetscan.org/vert_80/) (Agarwal et al., 2015). In order to be the more specific possible, we used a conservative approach, consisting of focusing only on genes presenting a 8-mer sequence in the 3’UTR region; hence with an exact match to positions 2-8 of the mature miRNA (the seed + position 8) followed by an ‘A’. The “enrichGO” function of the R package ClusterProfiler version 4.4.4 (Yu et al., 2012), with the GO dataset of the European sea bass was used to analyse function profiles of all genes potentially targeted by a given miRNA. A hypergeometric test was performed and enrichment p-value of gene ontology was calculated to find significantly enriched GO terms in the input list of each mirRNA target genes. Enrichment analysis was performed and a P value <0.05 was considered to indicate a statistically significant difference. The “ggplot 2” and “enrichplot” R packages were used to generate the cnetplot. The proportion of clusters in the pie chart was determined by the number of genes.

## 3. Results

### 3.1. Identification of miRNA differentially expressed between sex

Gonads of juveniles’ fish that were used for identification of sex markers were histologically differentiated, allowing to clearly discriminating males from females. Oocytes from ovarian tissues were at the primary growth stage (Fig. S1A), while some spermatocytes, spermatids and spermatozoids were distinguishable in the testis (Fig. S1B).

A total of 223 miRNAs, from the 10 samples sequenced (immature individuals), were annotated with *Prost!* (Desvignes et al., 2019). Among these miRNAs, 11 were differentially expressed between immature males and females (Table S2). Eight of these miRNAs were significantly more abundant in females, while the remaining three miRNAs were significantly more abundant in males. From that list of 11 miRNAs, we identified three miRNAs potentially acting on the regulation of genes involved in pathways of sexual development (based on the mammalian literature). Specifically, both miR-1388-3p and miR-7132a-5p were up-regulated in the females’ plasma (padj = 0.004 and p = 0.0003; Fig. 2A), while miR-499a-5p was more abundant in males’ plasma (padj = 0.0006, Fig. 2A). They were thus chosen for further validations by qPCR (Table 1). Both, miR-7132a-5p and miR-499a-5p were validated (Fig. 2B) on the sample sent to sequencing. To test for their consistency, additional immature individuals were analyzed (n = 10 on non-sequenced 1, n = 9 on non-sequenced 2; n = 27 sampled in the stress experiment, Table 1). No significant differences were observed between males and females, even though the miR-499a-5p tended to be higher in males than females for the three set of samples (Fig. S2). We also took advantage of the sequencing of individuals from the stress experiment to perform a DESeq2 analysis on the sex (all stressful conditions mixed; 9 M vs 11 F, Table 1). Only the miR-499-a-5p tended to be more abundant in plasma of males (adjusted p-value = 0.38, non-adjusted p-value = 0.01; Table S3).

**Fig. 2.**
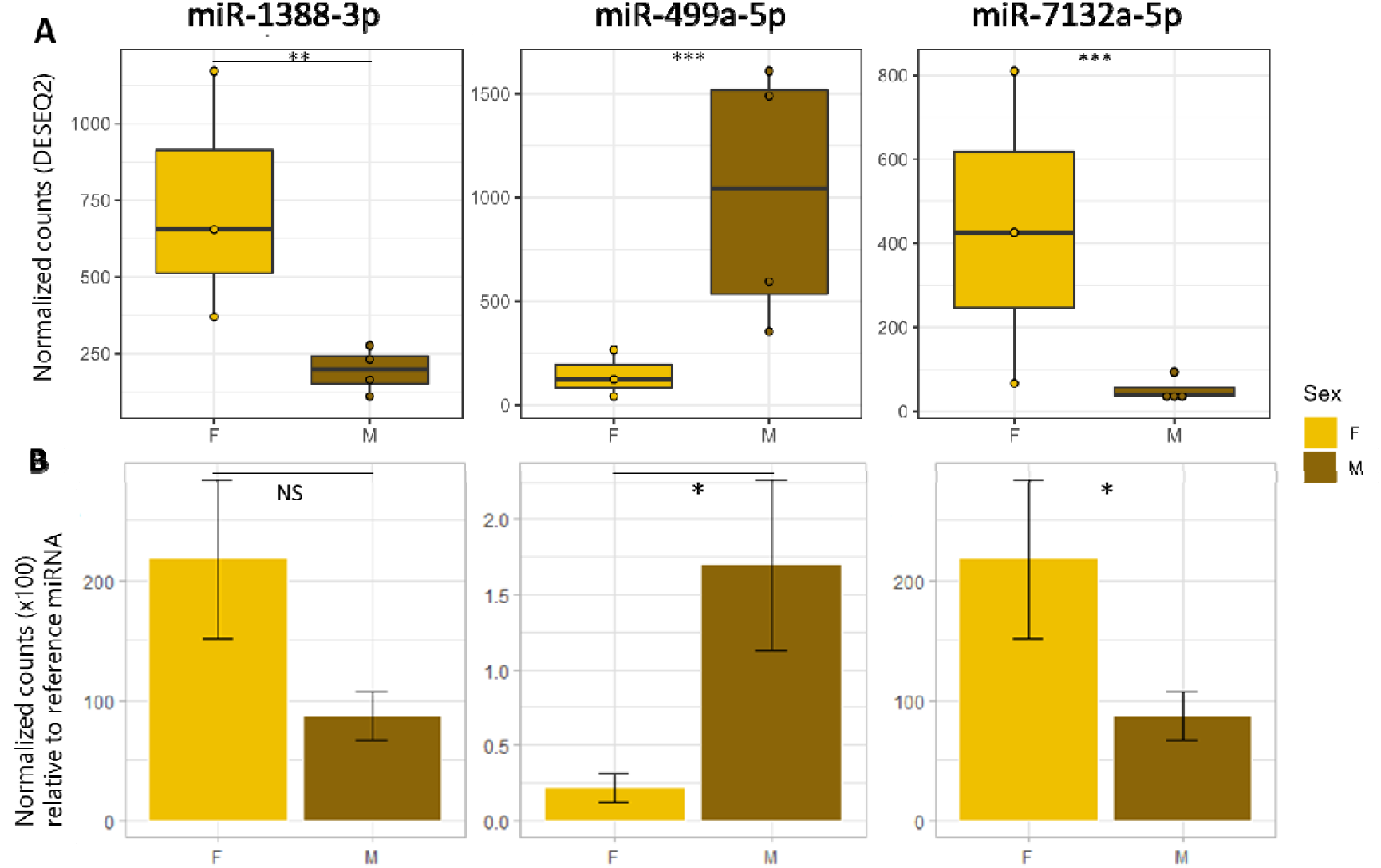
Normalized counts of circulating miRNAs based on A) small RNA-seq and B) RT-qPCR of sequenced samples. Reference miRNA used for RT-PCR relative quantification was the miR-23b-2-3p. NS: non-significant, *: p<0.05, **: p<0.01 and ***: p<0.001

A total of 441, 692 and 288 genes were predicted as potential target for respectively miR-1388-3p, miR–499a-5p and miR-7132a-5p (Table S4). For each miRNA, potential target genes were classified by their molecular function (MF) and biological processes (BP). On a total of 16368 possible biological processes in the European seabass, we detected only 59 significantly enriched GO for miR-7132a-5p, 103 significantly enriched GO for miR-499a-5p and 97 significantly enriched GO for miR-1388-3p (Table S5). On a total of 20503 possible molecular functions in the European seabass, we detected only 17 significantly enriched GO for miR-7132a-5p, 32 significantly enriched GO for miR-499a-5p and 28 significantly enriched GO for miR-1388-3p (Table S5). Within those GO of molecular functions and molecular process, many of them are involved in epigenetic regulation, mostly regarding histone methylation. Only the miR-499a-5p presented potential target genes in a GO directly related to sexual development: “Androgen receptor binding” (Table S5).

### 3.2. Dynamics of circulating miRNAs after an acute stress

The basal level of cortisol at T0 was 2.4 ±1.0 mg/ml (n=18; Fig. 3A). We observed a significant increase of cortisol at 0.5h (86.2 ±8.1 ng/ml, n=15, p-value = 1.4 x 10^-7^) and 1.5h (126.0 ±25.7 ng/ml, n=15, p-value = 9.7 x 10^-9^) following the confinement stress. Six hours after de stress, cortisol level decreased, though the fish did not return to their basal level (47.5 ±7.5ng/ml; Fig. 3A).

**Fig. 3.**
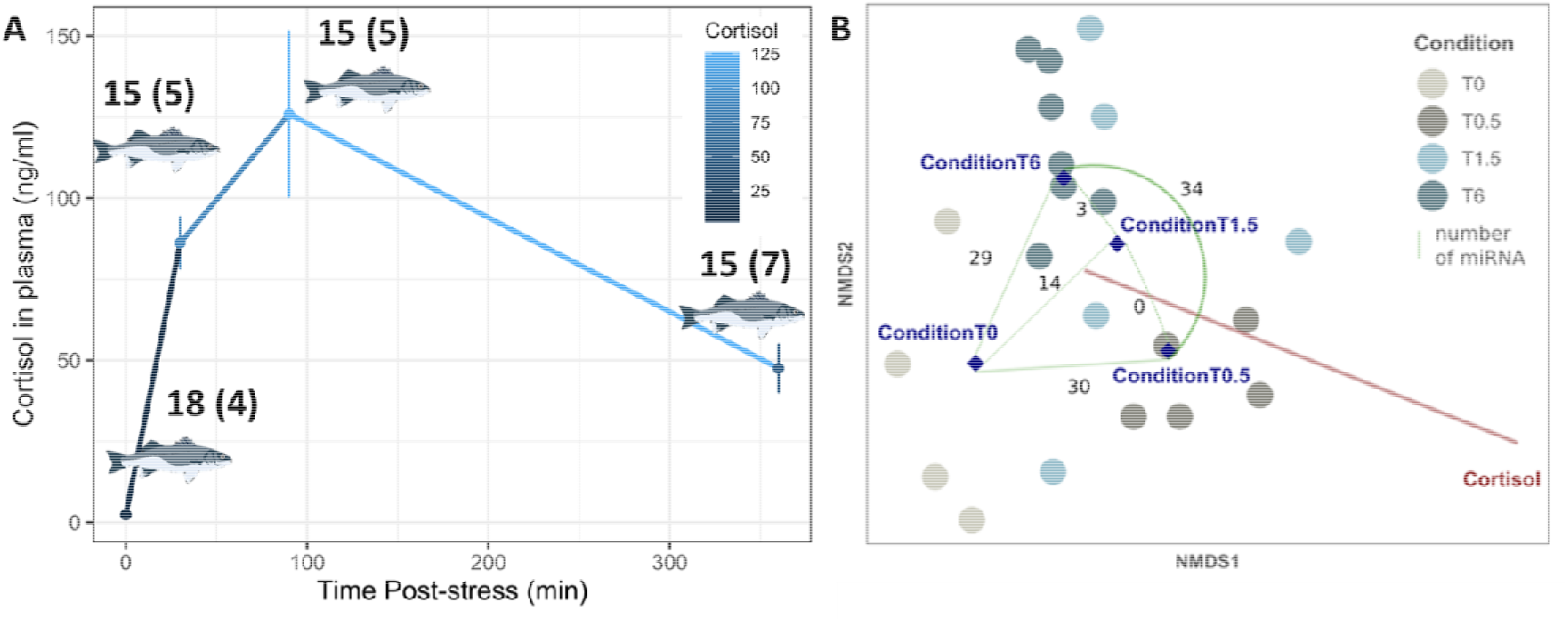
(A) Cortisol (ng/mL) quantified in plasma of European seabass after an acute stress of confinement and (B) Non-metric multidimensional scaling (NMDS) of miRNAs differentially expressed between various time points after confinement stress. Numbers in brackets corresponded to the number of samples used for sequencing. On NMDS, points represent individuals, diamonds represent the centroid value for each condition and the red line represent the fitted cortisol value.

A total of 257 miRNAs, from the 21 samples sequenced, were annotated with *Prost!*. Results from a 2-dimensional NMDS analysis yielded a stress level < 0.2 (stress = 0.11) and a noteworthy discrimination of control individuals from individuals of the T0.5 condition (Fig. 3B). The DESEQ2 analyses allowed to detect 30 miRNAs differentially expressed between T0 and T0.5; 14 miRNAs differentially expressed between T0 and T1.5; 29 miRNAs differentially expressed between T0 and T6 and 34 miRNAs differentially expressed between the T0.5 and the T6 conditions (Fig. 3B). Interestingly, the Venn diagram identified one miRNA (miR-200a-3p) as common for all comparison and was thus chosen for further qPCR validation (Fig. S3). We also purposely chose four other miRNAs from the above-described list. This choice was based on the fact that two of them: miR-155-5p and miR-205-1-5p (Fig. 4) followed the cortisol production dynamic, while another: miR-143-3p, presented an opposite pattern (Fig. 4). Indeed, the number of normalized counts between T0 and T0.5 significantly increased for miR-155-5p; miR-200a-3p and miR-205-1-5p whereas it decreased for the miR-143-3p (Fig. 4). Additionally, miR-223-3p was also selected because it presented positive linear correlation with cortisol production over time, and its expression was 2-fold higher at 6 hours post-stress compared to T0 (padj = 0.013; Fig. 4). Mir-155-5p, miR-200a-3p and miR-205-1-5p significantly increased in the 30 minutes following the confinement challenge (padj = 9,81.10^-8^; 5,65.10^-13^ and 9,03.10^-12^, respectively).

**Fig. 4.**
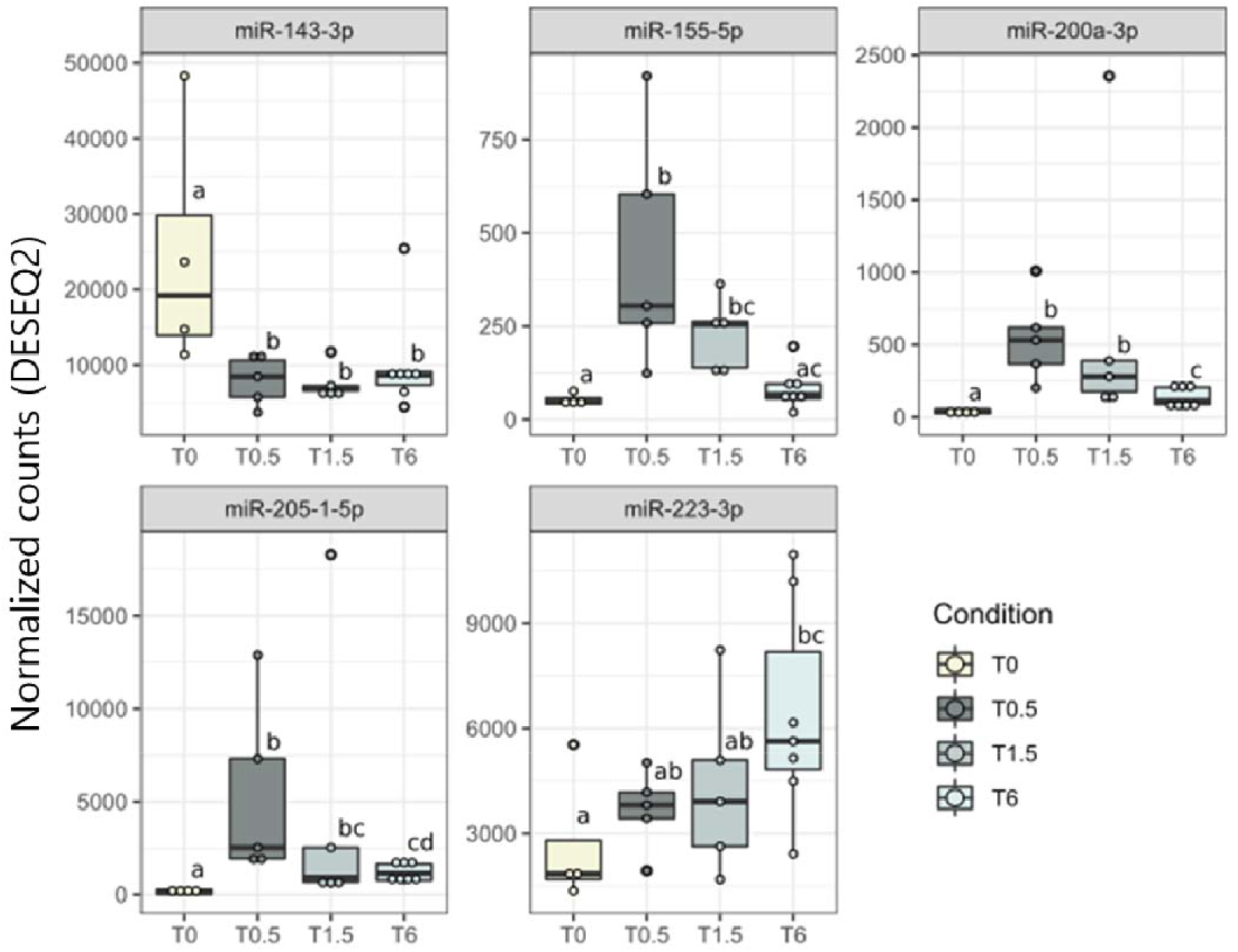
Normalized counts of miRNAs differentially expressed in plasma of European seabass after exposure to a confinement stress. Different letters reveal a significant difference. The color code is associated with timing of sampling after the acute stress.

The profile of expression observed with small RNA-sequencing and RT-qPCR was similar for miR-155-5p, miR-143-3p, miR-200a-3p and miR-223-3p but not for miR-205-1-5p (Fig. S4). To ensure that the miRNAs selected were effective markers, they were also tested by RT-qPCR but on other individuals (non-sequenced) from the same experiment. Regarding miR-143-3p, miR-200a-3p and miR-205-1-5p, they presented the same profile on RT-qPCR to that observed by sequencing (Fig. 5). This was especially true for miR-205-1-5p that increased significantly at 1.5 hour following the stressful challenge (p-value = 0.006). However, previous observations of the expression of miR-223-3p were not confirmed by RT-qPCR on those different samples (Fig. 5). In addition, the quantity of miR-155-5p was two low on these new samples for being correctly interpreted (i.e. very high CTs), and was thus discarded from the analysis.

**Fig. 5.**
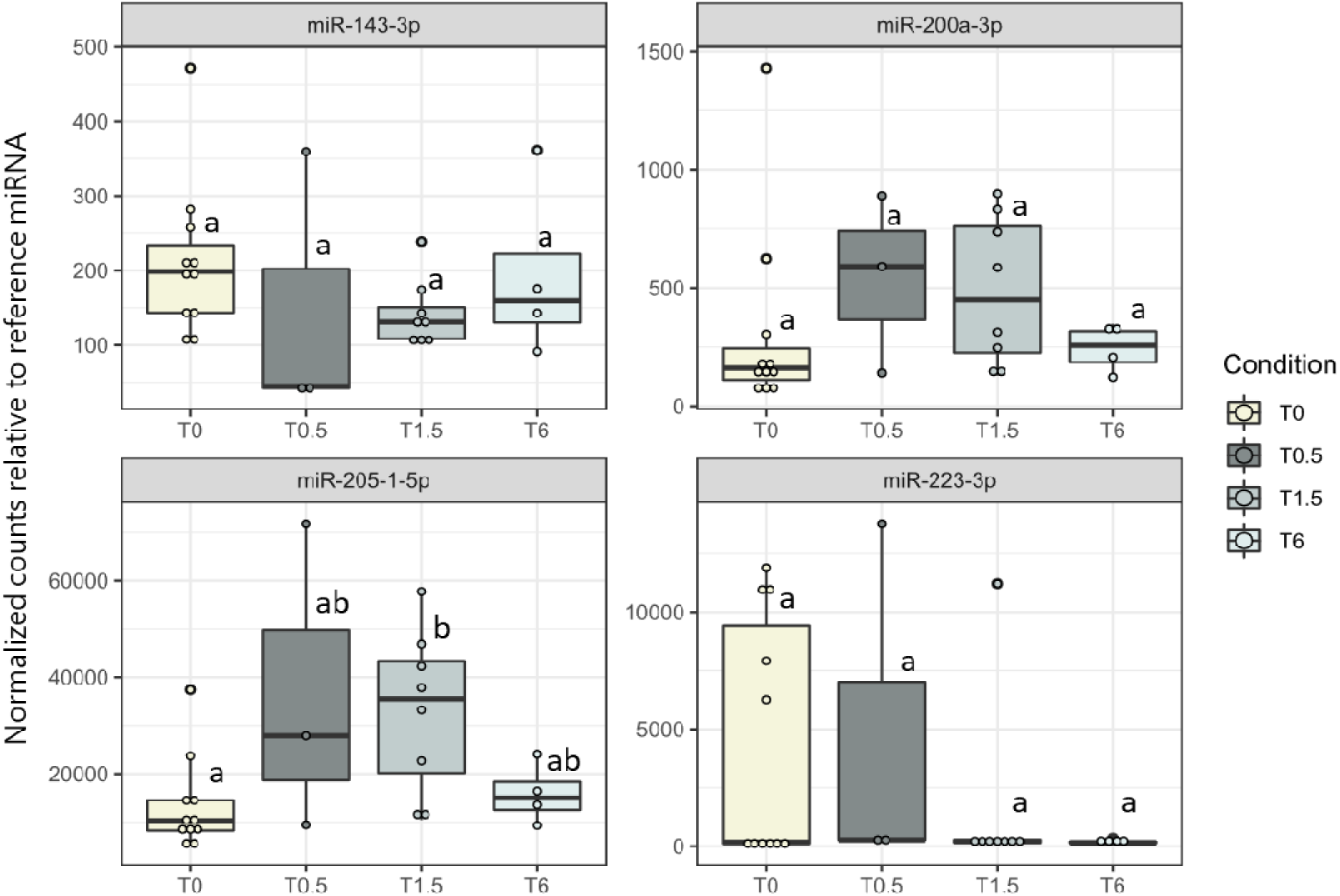
Relative expression profile of four miRNAs in plasma of European seabass after a confinement stress. RT-qPCR were done on non-sequenced samples collected at T0 (n=6), T0.5 (n=10), T1.5 (n=6) and T6 (n=8). Reference miRNAs used for RT-PCR relative quantification was miR-23b-2-3p. Different letter reveal a significant differences. The color code is associated with timing of sampling after the acute stress.

We focused only on those five miRNAs for the GO analysis. A total of 785, 739, 1146, 961 and 907 genes were predicted as potential target for respectively miR-155-5p, miR-143-3p, miR-200a-3p, miR-223-3p and miR-205-1-5p (Table S4). For each miRNA, potential target genes were classified by their molecular function (MF) and biological processes (BP). On a total of 16368 possible biological processes in the European seabass, we detected only 118 significantly enriched GO for miR-155-5p, 70 significantly enriched GO for miR-143-3p, 155 significantly enriched GO for miR-200a-3p, 116 significantly enriched GO for miR-223-3p and 138 significantly enriched GO for miR-205-1-5p (Table S6). On a total of 20503 possible molecular functions in the European seabass, we detected only 28 significantly enriched GO for miR-155-5p, 23 significantly enriched GO for miR-143-3p, 26 significantly enriched GO for miR-200a-3p, 33 significantly enriched GO for miR-223-3p and 19 significantly enriched GO for miR-205-1-5p (Table S6). Interestingly many pathways related to the primary (i.e. involving monoamine neurotransmitters and corticosteroids release) and secondary (i.e. metabolism, hydromineral balance and cardiovascular functions) response to stress were pinpointed in the GO analysis (Table S6, Fig. 6).

**Fig. 6.**
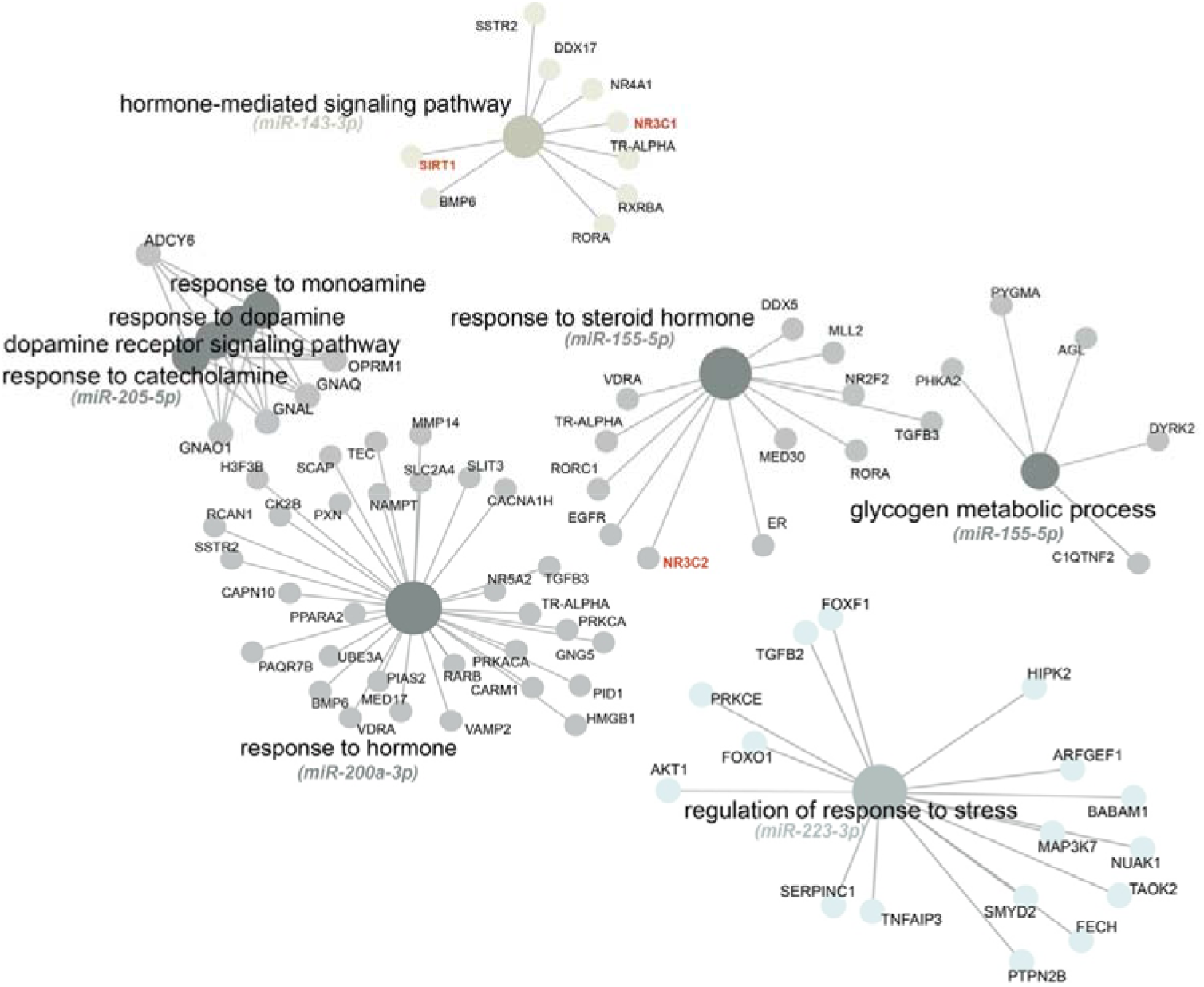
Key pathways of genes potentially targeted miR-143-3p, miR-205-1-5p, miR-155-5p, miR-200a-3p and miR-223-3p. The different colors are associated with the level of expression of each miRNA at each time point. See Fig. 4 and Fig. 5 for the description of colors code used.

For instance, within those GO of biological process, many of them are involved in the stress response: “hormone-mediated signaling pathways” (including key genes such as glucocorticoid receptor, nr3c1 and sirtuin, sirt1) for miR-143-3p; “response to steroid hormones” (including the mineralocortoid receptor, nr3c2) for miR-155-5p; “response to monoamines, catecholamine, dopamine” for miR-205-1-5p; “response to hormones” for miR-200a-3p and “regulation of response to stress” for miR-223-3p (Fig. 6). We also identified potential target genes involved in 1) behavioural response:”regulation of behaviour”, “social behaviour”; “regulation of locomotion” 2) regulation of blood pressure: “regulation of heart rate”, “angiogenesis”, “blood vessel development”, “heart development”; and 3) energy balance: “glycogen metabolic process”, “response to lipid”, “lipid modification”, “response to glucose”, “energy reserve metabolic process”, “response to insulin”, “negative regulation of protein metabolic process” (Table S6, see Fig. S5 for a detailed list of selected GO of biological processes, miRNA per miRNA).

## 4. Discussion

In this study, we investigated the potential of miRNAs as biomarker of sex and stress on the basis of a simple blood collection on the European seabass. We divided the research in two parts: searching for (i) sex markers and (ii) acute stress markers. Various miRNAs were modulated by the fact of being male or female, or by an acute stress.

Here, we identified eleven miRNAs interesting for sexing immature European seabass, three of them being strongly differentially expressed between sexes miR-1388-3p, miR-7132a-5p and miR-499a-5p. Among them, the miR-499a-5p, more detected in males, was likely the more interesting because of the similar profile obtained by RT-qPCR. In human, high level of circulating miR-499a is associated with early detection of breast cancer in women, while low levels were detected in cells of men with prostate cancer (Y. Chen et al., 2021; Kabirizadeh et al., 2016), suggesting a conserved role in steroid regulation. A recent study on zebrafish detected significantly more miR-499-5p in testis than in ovaries (van Gelderen et al., 2022). Yet, studies on other fish species have identified that miRNAs from the miR-499 family are involved in slow muscle phenotype determination (Duran et al., 2020) and the miR-499a-5p is key in regulating *Sox6* (Sex determining region Y-box 6) activity (Nachtigall et al., 2015; X. Wang et al., 2011). Interestingly, in the European seabass, the 3’UTR sequence of sox6 also have two possible 8-mer recognition sites for miR-499a-5p and *sox6* is significantly more expressed in the gonads of females experiencing molecular sex differentiation, compared to the gonads of males (Geffroy et al., 2021). This would support the possible repressive action of miR-499a-5p on *sox6*, though the exact mechanisms still remain to be established. Finally, the genes *kdm1a* (Lysine-specific histone demethylase 1A) and *prkcb* (Protein kinase C beta type) belonging to the GO “Androgen receptor binding” are also possible target for miR-499a-5p, supporting a possible role in sexual development. More studies are necessary to really depict the role of miR-499a-5p and understand why this miRNA is more detected in males’ plasma compared to females’ plasma.

Regarding miR-1388-3p (overexpressed in females of our experiment), this miRNAs was also more detected in ovary of *Paralichthys olivaceus* relative to testis (Yang et al., 2018). Furthermore, miR-1388-5p is involved in the regulation of the Spin-1 gene, which is important for hormone control, gametogenesis, and oocyte meiotic resumption in rainbow trout (F. Wang et al., 2020). Finally, the miR-7132a-5p (overexpressed in females of our experiment), was found to be up-regulated *Cynoglossus semilaevis* in pseudomales (ZW) compare to males (ZZ) (Zhao et al., 2021). In addition, the –3p strand of the miR-7132a was found to be up-regulated in the juveniles’ gonads of the common carp exposed to atrazine (F. Wang et al., 2019). It should be noted that none of the miRNAs tested in qPCR (in individuals that were not sequenced) were detected as significantly different between males and females, which might be due to the low quality of the RNAs used.

Regarding fish, our experiment corroborates observations of other studies conducted on gonads (miR-499-5p and miR-1388-5p), even though we were astonished to not identify other “usual suspects” involved in sexual development such as miR-202, which is highly sex-specific in the gonads of many other fish species (Geffroy et al., 2016; Juanchich et al., 2016; Qiu et al., 2018; Shen et al., 2023; J. Zhang et al., 2017) or statistically differentially expressed between the two gonads (Gay et al., 2018; J. Zhang et al., 2017). This might simply transduce that the miRNAs detected in the plasma does not specifically mirror what happen in other organs. In that sense, the recent study of Cardona and colleagues compared the miRNA repertoire of various body fluids or tissues of the rainbow trout and highlighted a large differences in the identity of the miRNAs observed in plasma compared to ovarian fluids (Cardona et al., 2021). Another explanation could be that miRNAs found in the plasma at a specific moment transduce a punctual physiological state at this moment, rather than a long-term phenotype (e.g., sex). This would explain why miRNAs related to stress were more reliably found in qPCR compared to those related to sex.

For the stress experiment, we first characterized the stress response of our fish, when exposed to confinement. This was highly challenging since the European seabass has often been regarded as a species with relatively high basal cortisol levels compared to other species (Alfonso et al., 2023). When considering individuals encompassing the same mass range (20-60 mg) the basal level varied between 14 and 80 ng/mL in other studies using a comparable ELISA quantification method (Cerqueira et al., 2020; Tsalafouta et al., 2015). Here, the level of cortisol was as low as 2 ng/ml, ensuring that we successfully obtained a reliable basal level of unstressed fish. To our knowledge, our study is the first to report a long term (6 hours) miRNA production dynamic following an acute stress in a fish species. A recent study described that 10 miRNAs were modulated one hour following an air exposure challenge in rainbow trout, in distinct non-lethal biological matrices (water, mucous and plasma; (Ikert et al., 2021). Here, the confinement stress affected the expression of 14 and 30 miRNAs at 0.5 and 1.5 hours post-stress, respectively. However, none of the miRNAs detected were similar to that of the Ikert et al. (2021) study. We could thus not tease out a specific effect of the stress applied or of the species studied, as pinpointed in a recent review (Raza et al., 2022). For instance, miR-210-3p has been associated with hypoxic stress in the rainbow trout (Cardona et al., 2022) and miR-276b-3p was shown to be upregulated following a salinity stress challenge in *Portunus trituberculatus* (X. Chen et al., 2019). Among miRNAs that we observed differentially expressed during stress recovery, we purposely chosen to focus on miR-155-5p, miR-143-3p, miR-200a-3p, miR-223-3p and miR-205-1-5p to conduct the RT-qPCR validation part. This choice was mainly based on their potentially interesting targets (discovered thanks to TargetScan) following a stress. At the physiological level, the stress response is conducted by the central nervous system, and led to a secretion of neurotransmitters and stress hormones that constitute the first stress response (Schreck & Tort, 2016). The second response involve the secretion of energetic metabolites (glucose, lactate..), the modulation of osmoregulation and immune response, while the third response lead to drastic changes in performances (decrease of growth, disease resistance, behavioral change..). Interestingly, the GO analysis allowed us to provide some hypothesizes regarding the down-regulation of key genes involved in the first two stages of the stress response. For instance, we identified the glucocorticoid receptor (*gr* or *nr3c1*) as potential target of miR-143-3p *in silico.* Such a link would make sense since 1) miR-143-3p is highly expressed at T0 compared to other time-points, supporting a possible production of GR post-stress and 2) gain– and loss-function approaches in GR and miR-143-3p confirmed that the glucocorticoid receptor was indeed a target of miR-143-3p in humans (L. Zhang et al., 2020). This support a specific role of miR-143-3p in the primary stress response. MiR-155-5p and miR-200a-5p, observed in the mid-intestine, are mainly involved in the response to an oxidative stress of the Wuchang bream*, Megalobrama amblycephala* (Song et al., 2021). It is worth noting that the mineralocorticoid receptor (mr or nr3c2), a potent receptor of cortisol (Prunet et al., 2006), is a potential target of miR-155-5p in our *in silico* analyses, supporting its possible role in the regulation of stress response. The miR-223 was also associated to the modulation of oxidative stress response in the Nile tilapia (Tang et al., 2013). We observed that the miR-205-1-5p was the only one to show a statistical and steadily increase of its relative expression following the confinement stress. Our predictive analysis, using TargetScan Human, allowed to detect several pathways related to stress such as “regulation of response to stress”, energy balance, such as “response to insulin”, “lipid modification” as well as “angiogenesis” and “negative regulation of blood circulation”. In human model, miR-205-5p regulates the VEGFA-angiogenesis (Oltra et al., 2020), supporting a role in the secondary stress response. In fish studies, miR-205-5p was also detected following stressful events like heat stress or hypoxia (Lai et al., 2016; Liu et al., 2022).

Challenges reported in diagnostic of human cancer are linked to technical issues and individual-related parameters that could influence the presence/absence of circulating miRNA (Tiberio et al., 2015). In this review, authors reported the various technical parameters (hemolysis, anticoagulant used, extraction method, miRNA measurement, data normalization) that could explain the differences in studies outcomes. Here, we also detected many limitations that are likely explained by differences in the quality of the samples, since for those of high-quality, small RNA-seq was confirmed by qPCR, but not for samples of apparently lower quality. Finally, a major difference between our study and the vast majority of the above-mentioned studies is that most of the other authors worked on tissues to identify the modulation of miRNA content in response to a stress or a physiological status. In fishes, miRNAs have been identified in several matrices, including non-lethal ones such as plasma, mucus, water and feces. Here we focused on plasma, that directly reflect the physiological state of an individual and this opens new perspectives of use and application to follow natural and captive livestock. Following the stressful challenge, we identified several key miRNAs that are readily released in blood (increasing in the 30 min after stress event) and return to their basal level in few hours. However, the information gathered highlighted that this method is likely too precise to provide cues on the sex, as it will rather indicates a physiological state linked to the gonadal development stage.

## Acknowledgments

The authors would like to acknowledge the team from Station Ifremer Palavas-les-Flots (France) and the team from Aquanord Gloria Maris (France) for the help in different sampling. The authors also thank Pierre Lopez for the sea bass infographic.

## Data accessibility

All data generated or analyzed during this study are included in this article and supplementary information.

## Author Contribution

Conceptualization, Benjamin Geffroy, Camille Houdelet; Methodology: Benjamin Geffroy, Camille Houdelet, Sophie Hermet, François Ruelle, Gilbert Dutto, Aline Bajek; Formal Analysis, Eva Blondeau-Bidet, Mathilde Estevez-Villar, Xavier Mialhe; Writing original draft, Camille Houdelet, Benjamin Geffroy; Writing-Review and Editing, Benjamin Geffroy, Camille Houdelet, Eva Blondeau-Bidet and Julien Bobe, Funding Acquisition, Benjamin Geffroy. All authors reviewed the manuscript.

---

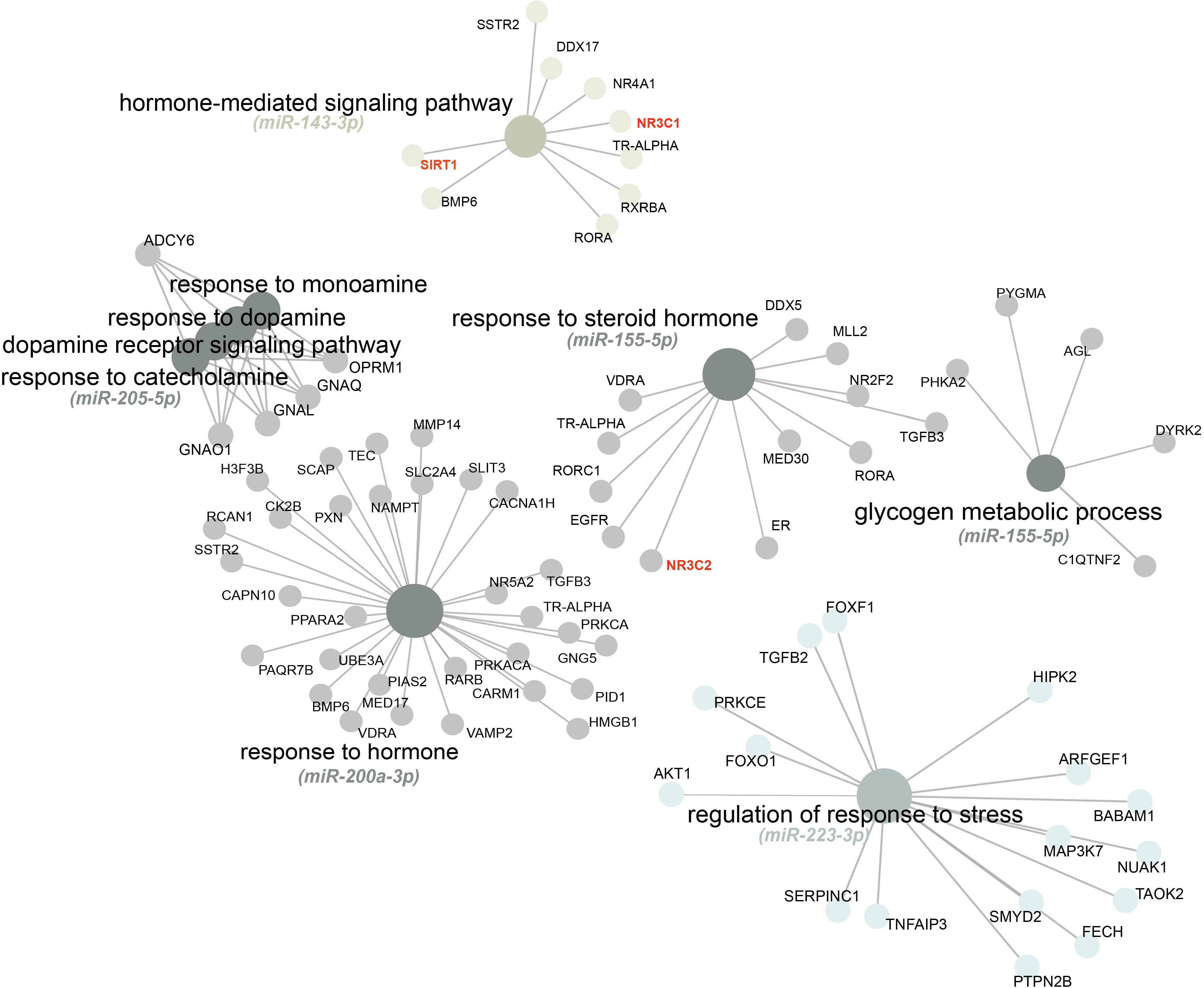

