## Supplementary material for "Circulating MicroRNAs indicative of sex and stress in the European seabass (*Dicentrarchus labrax*): toward the identification of new biomarkers": Figure S1

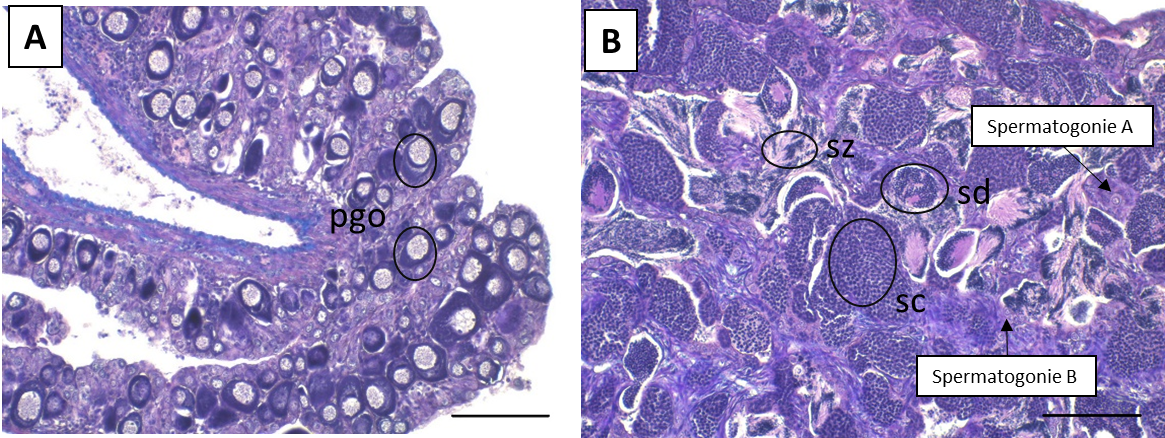

**Figure S1:** Histological sex of juveniles’ European seabass (x20). (A) Differentiated female gonad represented by primary growth oocytes (pgo). (B) Differentiated male gonad represented by spermatids (sd), spermatocytes (sc) and spermatozoa (sz). Scale: 100 µm.
