## Supplementary material for "Circulating MicroRNAs indicative of sex and stress in the European seabass (*Dicentrarchus labrax*): toward the identification of new biomarkers": Figure S2

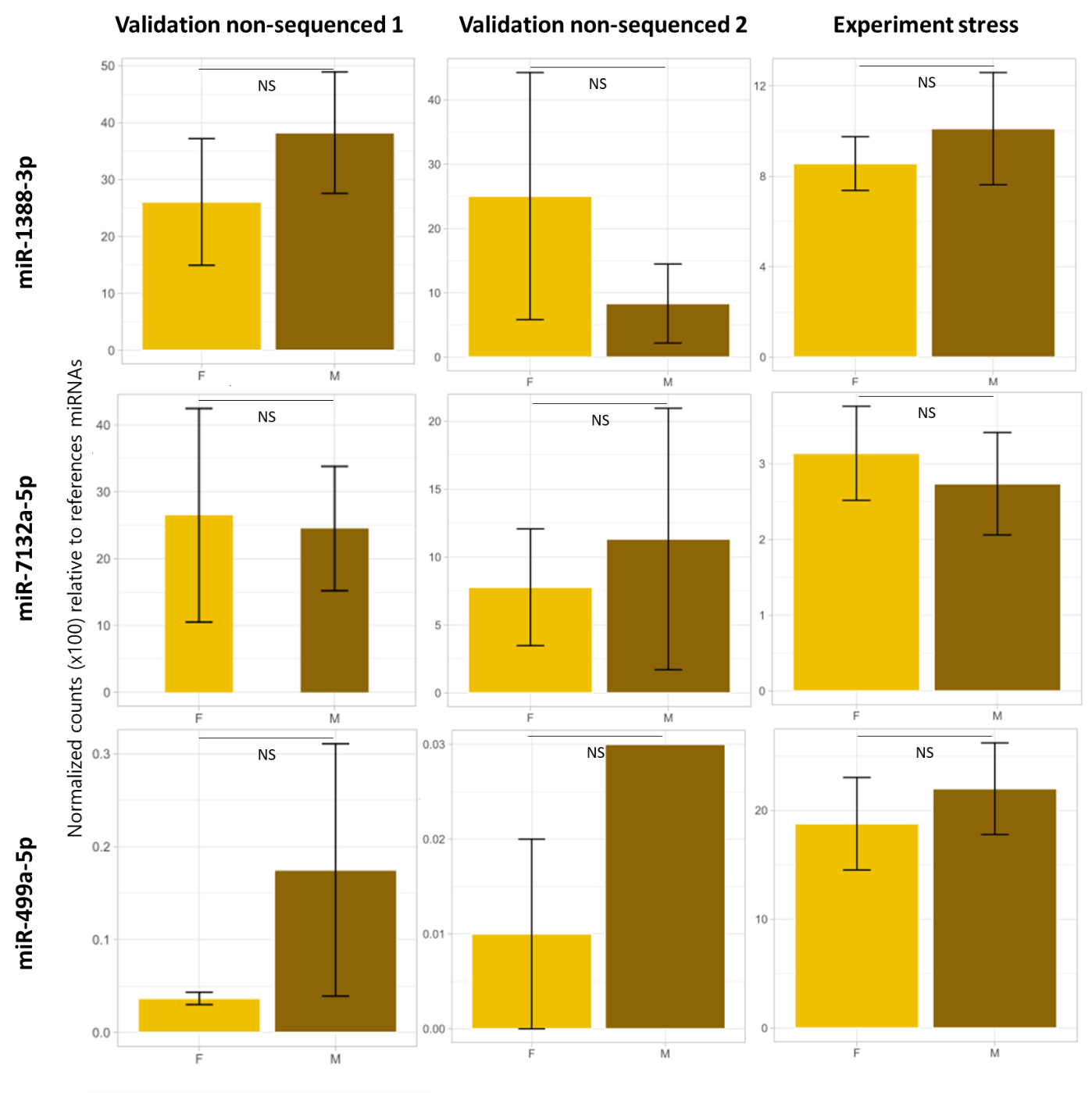

**Fig. S2** RT-qPCR expression of differentially expressed miRNA of non-sequenced samples from three independent experiments. Mean expression of miR-7132a-5p, miR-1388-3p and miR-499a-5p. Reference miRNA used for RT-PCR
