## Supplementary material for "Circulating MicroRNAs indicative of sex and stress in the European seabass (*Dicentrarchus labrax*): toward the identification of new biomarkers": Figure S3

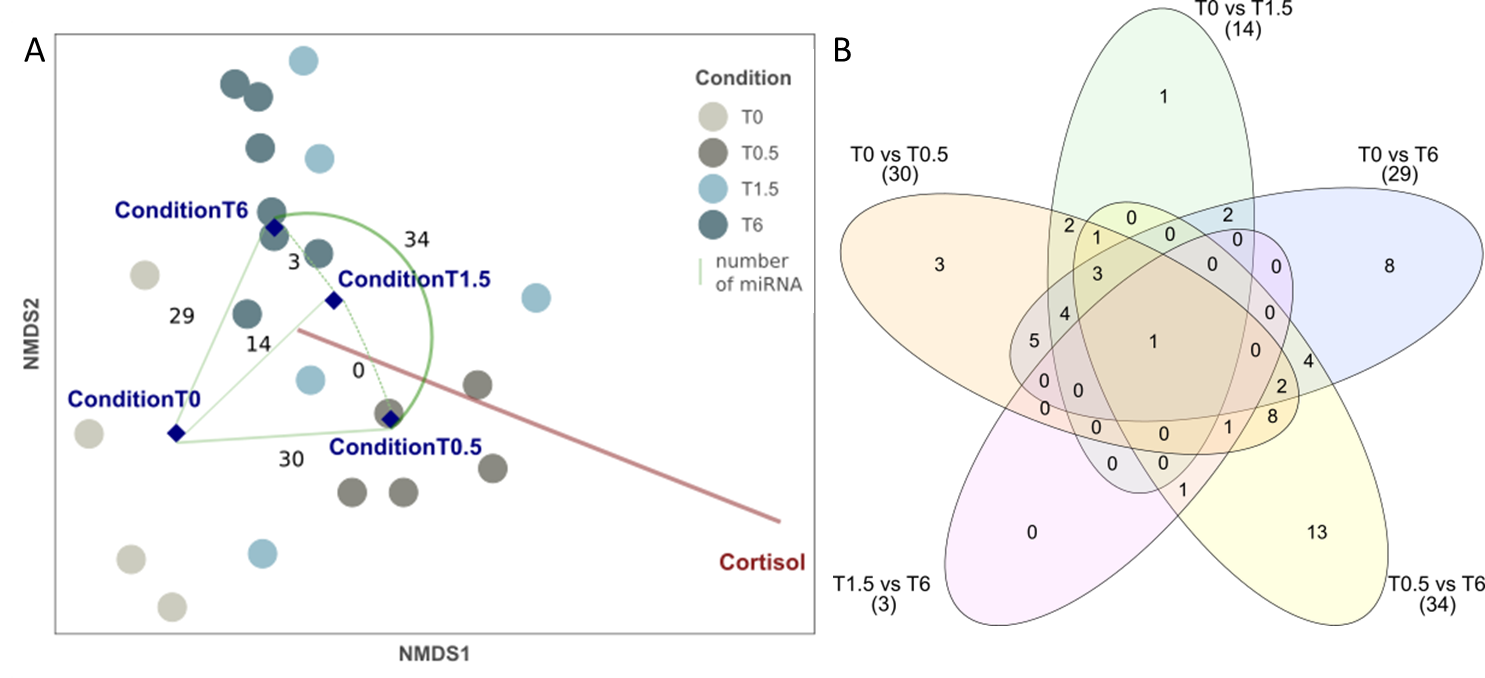

**Figure S3:** Venn diagram of miRNA differentially expressed between various time point after confinement stress.
