## Supplementary material for "Circulating MicroRNAs indicative of sex and stress in the European seabass (*Dicentrarchus labrax*): toward the identification of new biomarkers": Figure S4

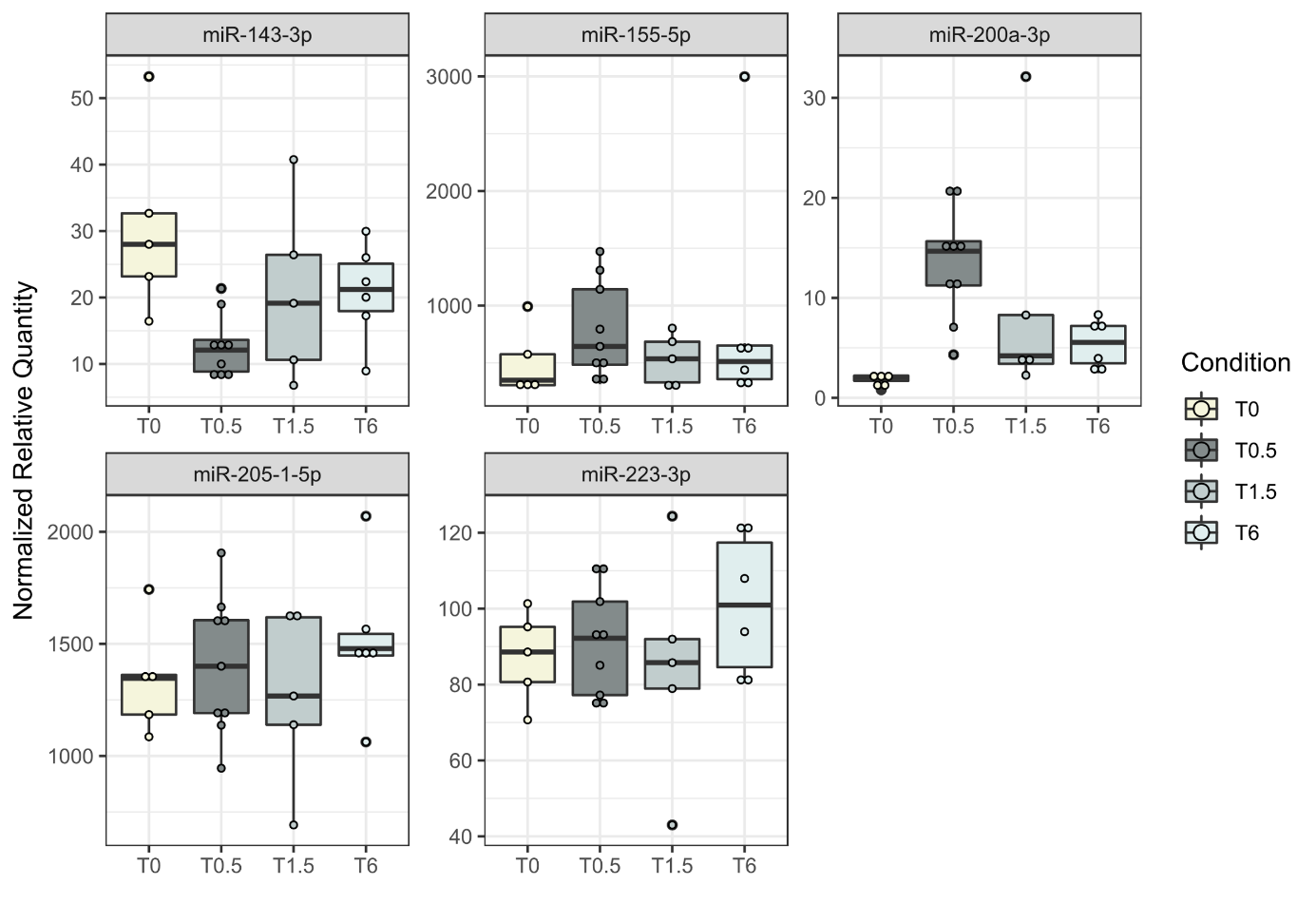

**Figure S4** RT-qPCR expression of differentially expressed miRNA of sequenced samples after a confinement stress. References miRNA used for RT-PCR relative quantification was miR-23b-2-3p. NS: non-significant.
