## Supplementary material for "Circulating MicroRNAs indicative of sex and stress in the European seabass (*Dicentrarchus labrax*): toward the identification of new biomarkers": Figure S5

### Slide 1
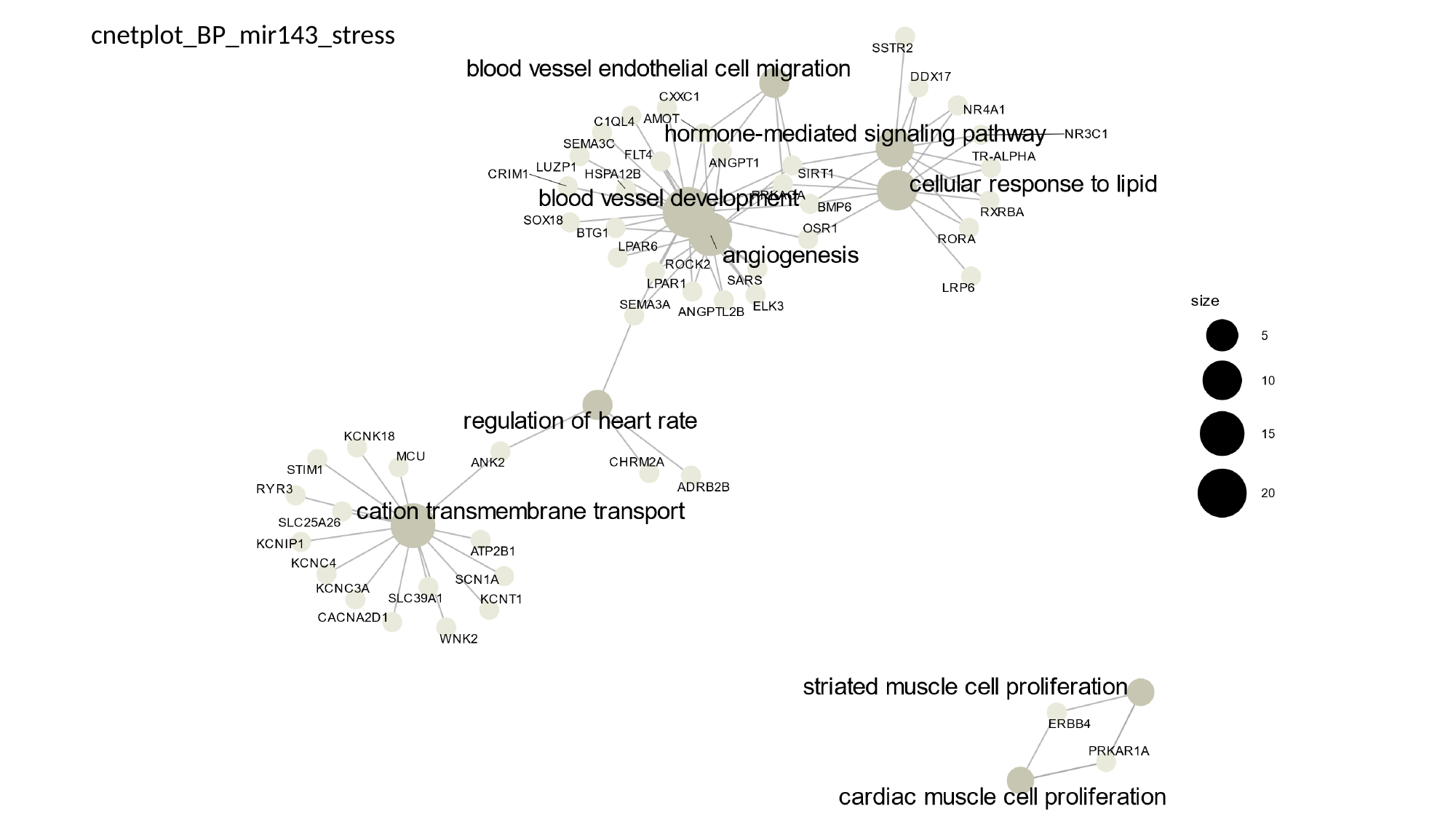

cnetplot_BP_mir143_stress

### Slide 2
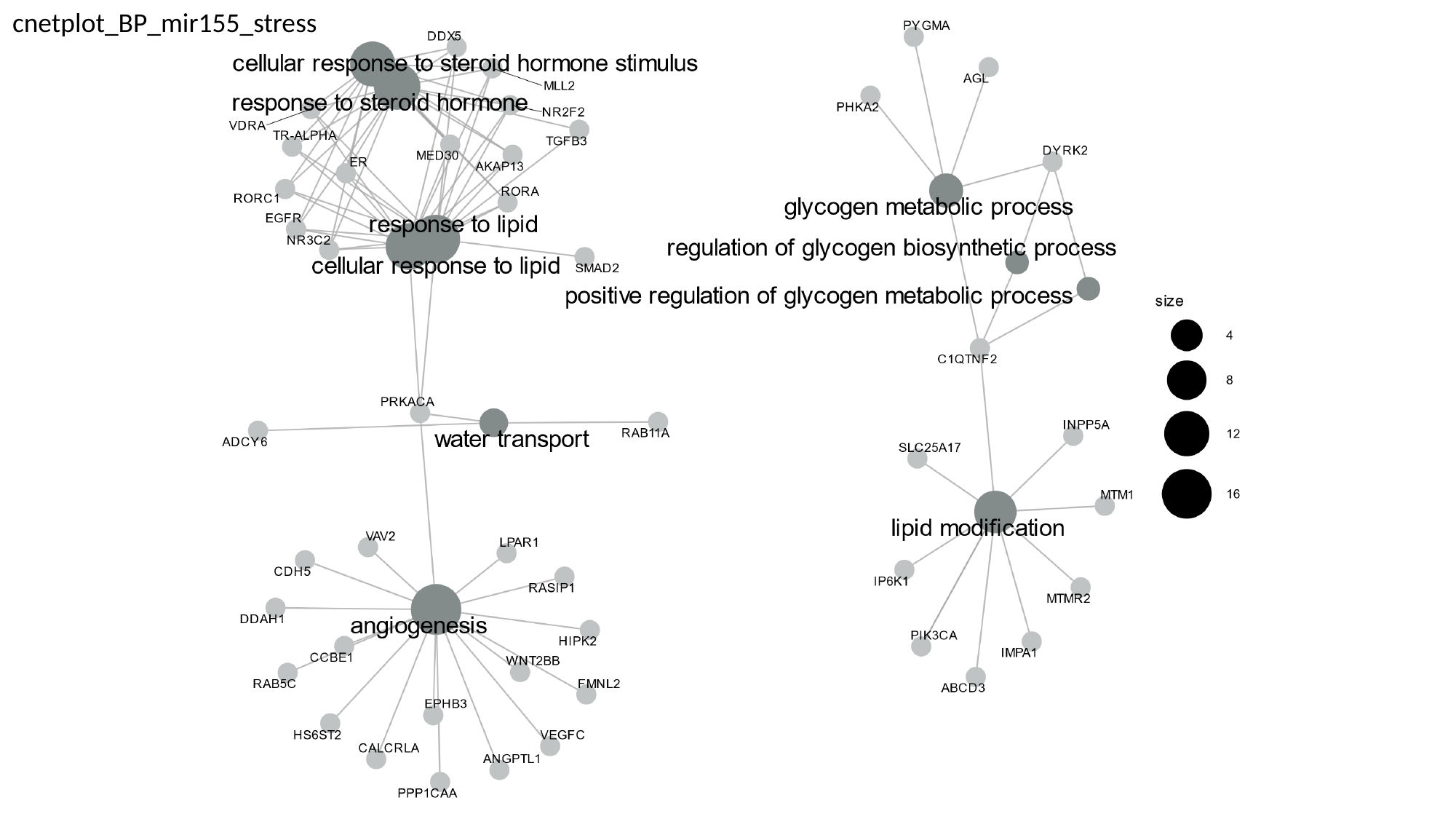

cnetplot_BP_mir155_stress

### Slide 3
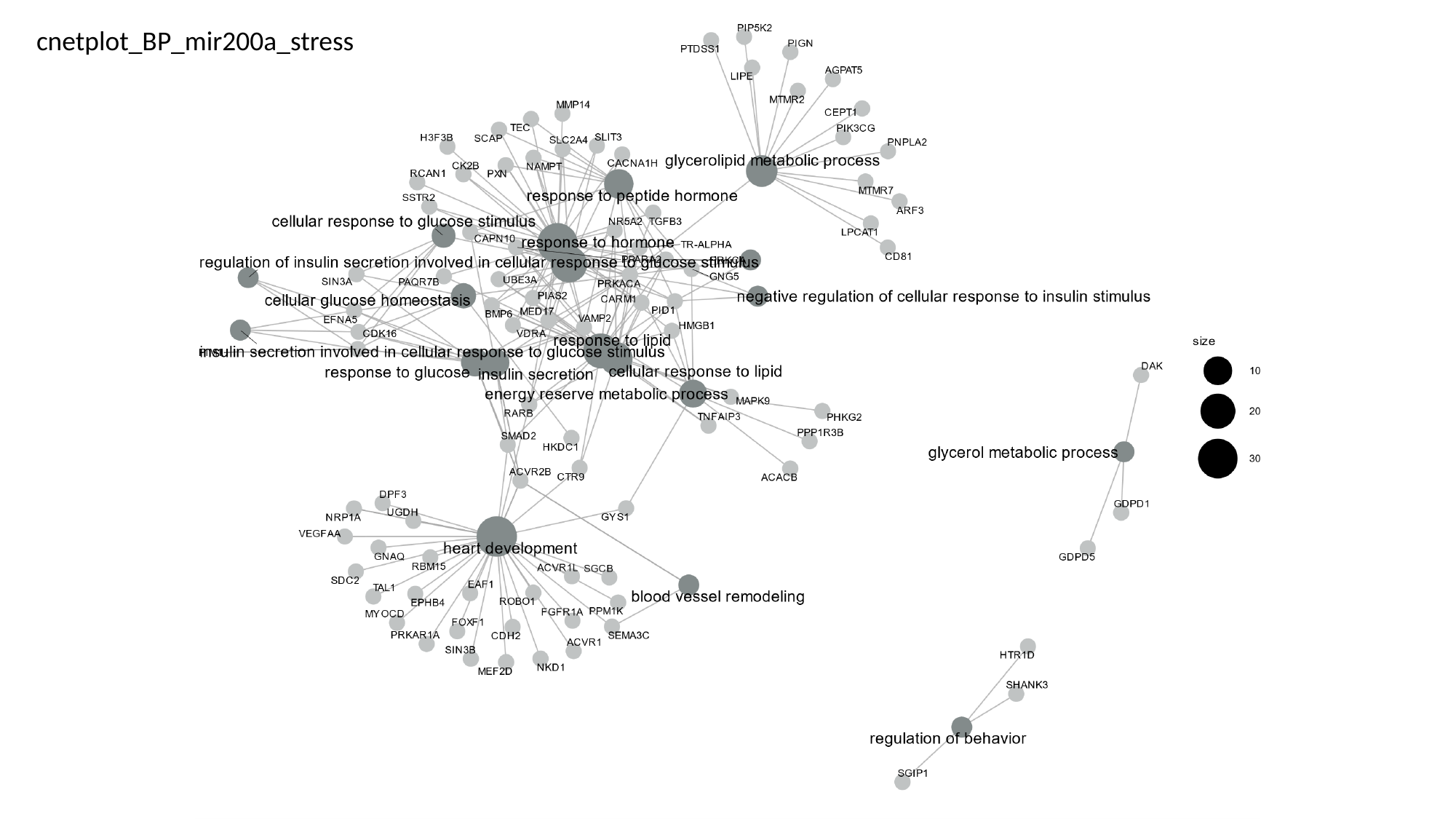

cnetplot_BP_mir200a_stress

### Slide 4
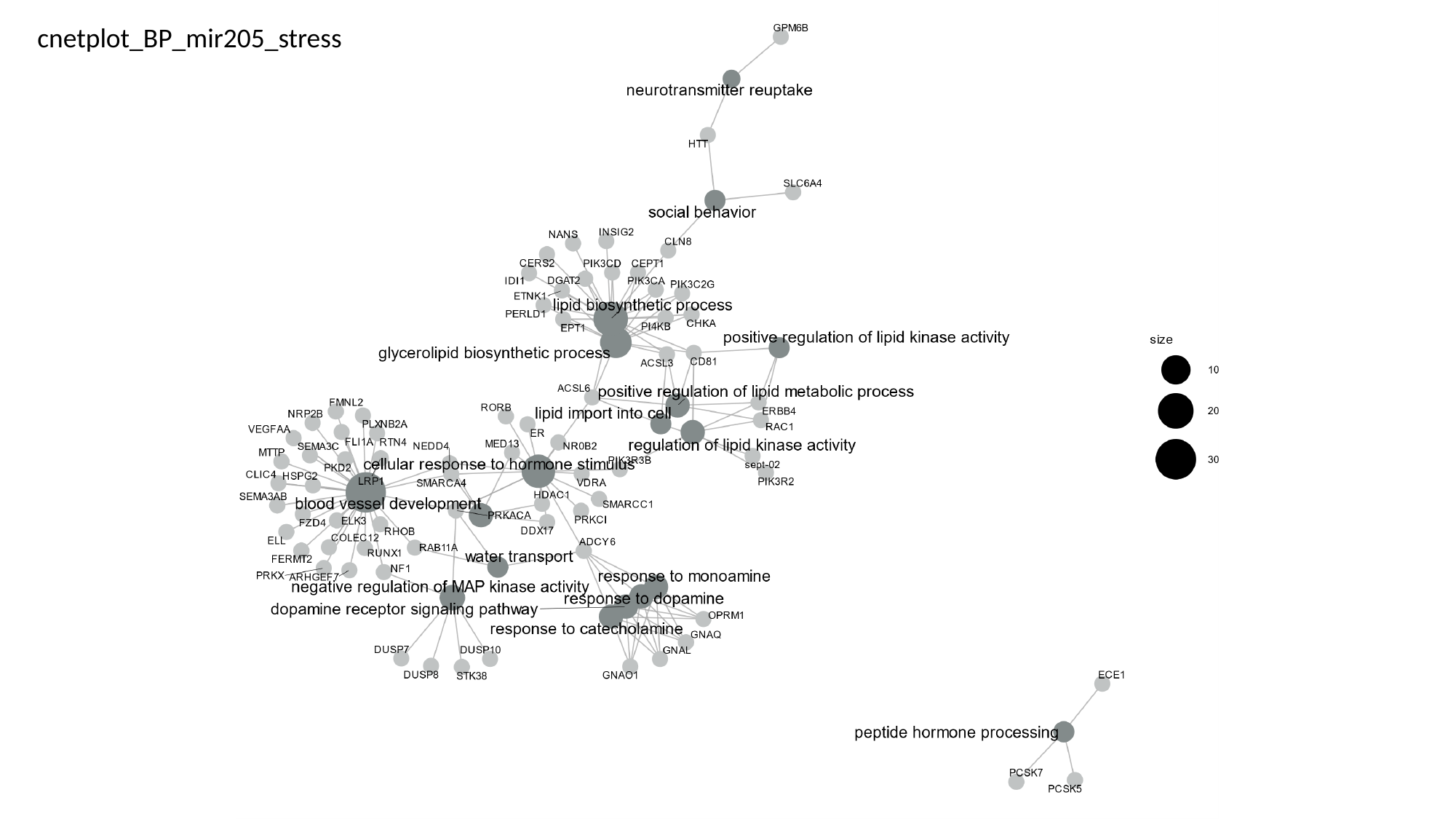

cnetplot_BP_mir205_stress

### Slide 5
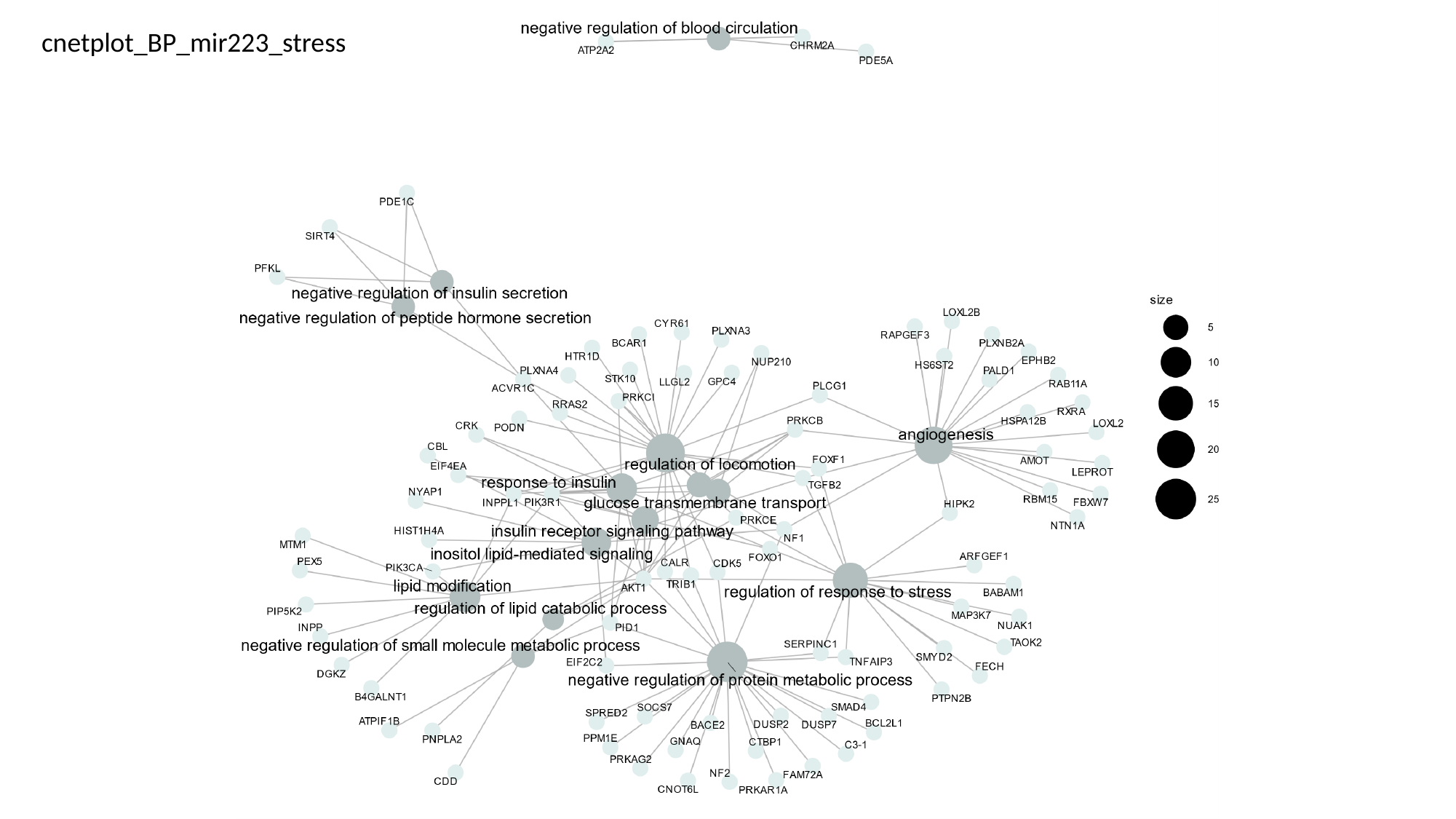

cnetplot_BP_mir223_stress
