## Supplementary material for "Circulating MicroRNAs indicative of sex and stress in the European seabass (*Dicentrarchus labrax*): toward the identification of new biomarkers": Table S2

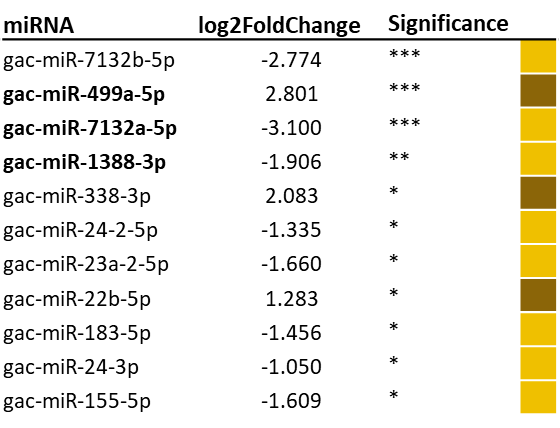

**Table S2:** list of miRNA differentially expressed between males and females (DESeq2). *: p<0.05, **: p<0.01 and ***: p<0.001. Yellow or brown boxes revealed miRNA mostly expressed in females or males, respectively.
